# Joint RElaxation-Diffusion Imaging Moments (REDIM) to probe neurite microstructure

**DOI:** 10.1101/598375

**Authors:** Lipeng Ning, Borjan Gagoski, Filip Szczepankiewicz, Carl-Fredrik Westin, Yogesh Rathi

**Affiliations:** Brigham and Women’s Hospital, Harvard Medical School, Boston, MA; Boston Children’s Hospital, Harvard Medical School, Boston, MA

## Abstract

Joint relaxation-diffusion measurements can provide new insight about the tissue microstructural properties. Most recent methods have focused on inverting the Laplace transform to recover the joint distribution of relaxation-diffusion. However, as is well-known, this problem is notoriously ill-posed and numerically unstable. In this work, we address this issue by directly computing the joint moments of transverse relaxation rate and diffusivity, which can be robustly estimated. To zoom into different parts of the joint distribution, we further enhance our method by applying multiplicative filters to the joint probability density function of relaxation and diffusion and compute the corresponding moments. We propose an approach to use these moments to compute several novel scalar indices to characterize specific properties of the underlying tissue microstructure. Furthermore, for the first time, we propose an algorithm to estimate diffusion signals that are independent of echo time based on the moments of the marginal probability density function of diffusion. We demonstrate its utility in extracting tissue information not contaminated with multiple intra-voxel relaxation rates. We compare the performance of four types of filters that zoom into tissue components with different relaxation and diffusion properties and demonstrate it on an in-vivo human dataset. Experimental results show that these filters are able to characterize heterogeneous tissue microstructure. Moreover, the filtered diffusion signals are also able to distinguish fiber bundles with similar orientations but different relaxation rates. The proposed method thus allows to characterize the neural microstructure information in a robust and unique manner not possible using existing techniques.

## I. INTRODUCTION

Magnetic resonance imaging (MRI) modalities, such as diffusion MRI (dMRI) and T_2_ relaxometry, probe different physical or biological properties of brain tissue. For example, T_2_ relaxation time is inherently related to the biochemical composition of the tissue (e.g. myelin), while dMRI is sensitive to the tissue meso and microstructure. The standard analysis approach in dMRI is based on imaging data acquired at a fixed echo time (TE) and assumes that the underlying diffusion is independent of T_2_ relaxation time. Joint analysis of dMRI data acquired at different TE allows to characterize the coupling between T_2_ relaxation and diffusion and provides unique information about the tissue properties that are not available using the standard methods. Consequently, multidimensional correlation of transverse relaxation rates and diffusion coefficient of water molecules has been recently investigated to understand tissue microstructure [1], [2], [3], [4], [5], [6], [7]. Diffusion-relaxation correlation spectroscopy [8], [9] is a classical approach to estimate the joint probability density function (PDF) of relaxation and diffusion. In this approach, the joint PDF of relaxation and diffusion is estimated by numerically solving an inverse Laplace transform [8], [10], [9] using a large number of imaging measurements acquired with different combinations of b-values and TE. A well-known limitation of this approach is that the estimation of the inverse Laplace transform is highly ill-posed [11]. As a result, several methods have been proposed to improve its numerical stability by adding suitable constraints in the computational algorithms or using information from neighboring voxels [12], [1], [2], [13]. While these methods help improve the estimation, yet they still require a large number of imaging measurements, and it is not clear if the obtained joint PDF is accurate in all scenarios. This further limits their application in clinical settings.

In this work, we address these challenges and propose a novel approach to characterize tissue microstructural properties without explicitly solving the inverse Laplace transform. Below, we list several distinctive features of our approach and the novel contributions of this work.

1. Our approach directly estimates the joint moments of the relaxation and diffusion using the standard least-squares algorithm which is numerically more reliable than estimating the inverse Laplace transform.
2. Our approach includes a novel algorithm that uses several multiplicative filters to rescale the joint PDF of relaxation-diffusion in order to characterize (or “zoom-into”) more specific properties of the underlying joint distribution of relaxation and diffusion.
3. We use the joint moments of the rescaled joint PDFs to estimate the principal diffusion directions associated with different transverse relaxation rates.
4. Experimental results on in-vivo human brain data show that, in fiber crossing areas, different fiber bundles could potentially have different transverse relaxation rates. For the first time, we demonstrate that dMRI signals based on filtered PDFs are able to separate fiber bundles with different relaxation rates, which has important implications for diffusion tractography.

Thus, the proposed method provides novel information about the tissue meso and microstructure and reveals the heterogeneous relaxation-rates in different axonal fiber bundles.

## II. THEORY

For a water molecule diffusing in a medium with traversal relaxation rate *r* = 1*/T*_2_ and the diffusivity along a direction ***u*** of *D*(***u***), it is standard to model the decay of dMRI signal due to relaxation and diffusion using the exponential functions exp(*−rt*) and exp(*−bD*(***u***)), where *t* and *b* denote the TE and b-value. Therefore, the dMRI signal from water molecules with different relaxation and diffusion can be modeled jointly by

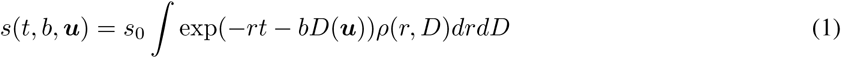

where *ρ*(*r, D*) represents the joint probability distribution of *r* and *D*. Since relaxation and diffusion reflect different properties of the underlying tissue, the statistical properties of *ρ*(*r, D*) are important to characterize tissue microstructure.

The model in Eq. (1) provides a general expression for dMRI signals. In the special case when *t* is fixed, (1) becomes a model for standard multi-shell dMRI signal based on diffusion tensor distributions as in [14], [15]. It is able to model the non-exponential decay of dMRI signals with increasing b-values. It also includes the standard multi-exponential (and multi-compartmental) models [16], [17], [18] as special cases. In the case of multiple TE, the signal model Eq. (1) has been considered in [8], [2].

Eq. (1) implies that the joint PDF *ρ*(*r, D*) can be estimated via the inverse Laplace transform of the normalized signal *s*(*t, b*)*/s*_0_. But this approach not only requires long scan time but also is extremely ill-posed and hence numerically unstable. On the other hand, one can reliably estimate the moments of this joint distribution, which can be used to derive several novel quantitative measures that provide new information about the tissue microstructure as described in the following sections. Next, we introduce an algorithm to estimate the joint moments of *ρ*(*r, D*) and an approach to derive novel measures based on these moments.

### A. On cumulant expansion and moment estimation

Cumulant expansion is a standard technique in dMRI to approximate the signal using the lower order moments of the diffusion propagator. The diffusion tensor model is a special case when only the second order moment tensor is considered. The diffusion kurtosis imaging technique involves both the second and fourth order moment tensors. The cumulant expansions have also been applied to estimate water exchange using dMRI experiments [19]. Below, we adapt this technique to derive the cumulant expansion of *s*(*t, b*, ***u***).

We let ⟨*r⟩*_*ρ*_ and ⟨*D*(***u***)⟩_*ρ*_ denote the expected value of *r* and *D*(***u***) respectively. Moreover, we denote

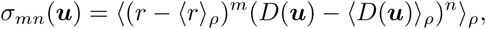

as the (*m, n*)^*th*^ central moments (*m >* 0, *n >* 0) of the joint distribution of relaxation and diffusivity along a gradient direction ***u*** but where the relaxation rate *r* is assumed to be independent of the gradient directions. Next, we introduce the following cumulant expansion of the diffusion signal:

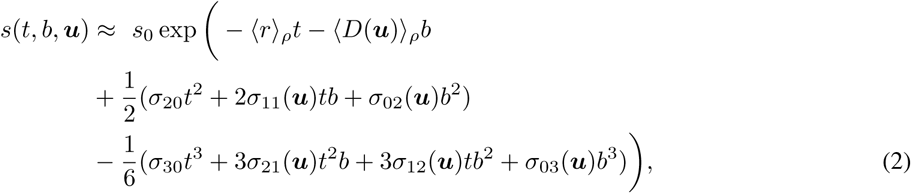

where ⟨*r*⟩_*ρ*_, *σ*_20_, *σ*_30_ are independent of ***u***. In the above expression, we use the third-order expansion to explore information beyond the DKI technique.

By taking the logarithm of both sides of Eq. (2), we obtain

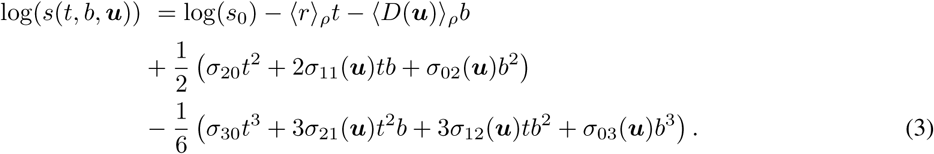

In above, *t* and *b* are known parameters determined by the dMRI sequence. The unknown parameters ⟨*r*⟩_*ρ*_, ⟨*D*(*u*))⟩_*ρ*_ and *σ*_*mn*_(***u***) can be estimated using the standard least-squares algorithm by jointly fitting diffusion signals along different directions using Eq. (3). A total number of 9 parameters are obtained along each gradient direction. We note that ⟨*D*(*u*))⟩_*ρ*_ is not restricted to be a quadratic function as in DTI. Thus, the model is suitable for analyzing signals from tissue with crossing fibers. Moreover, the signals along different directions are coupled via the parameters ⟨*r*⟩_*ρ*_, *σ*_20_, *σ*_30_.

### B. Moments of filtered relaxation-diffusion probability density functions

Based on the estimated model parameters in Eq. (3), it is straightforward to compute the joint moment ⟨*r*^*m*^*D*(***u***)^*n*^⟩_*ρ*_. For example

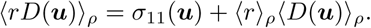

Generally, physically meaningful summary indices, such as mean kurtosis [20], are used to characterize the statistical properties of high dimensional variables. Consequently, we propose an approach to derive several summary indices based on the joint moments of *r* and *D*(***u***). To simplify notations, we derive expressions for *D* along a particular direction ***u***, thereby dropping the variable ***u***. However, we should note that, the obtained expressions are valid for any ***u*** and hence directionally sensitive.

For a nonnegative function *f* (*r, D*), the expected value under a given probability distribution function *ρ*(*r, D*) is given by

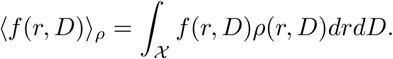

The function *f* (*r, D*) is a multiplicative filter that rescales the joint PDF *ρ*(*r, D*) in order to emphasize the signal from a particular tissue component. Then the rescaled PDF *ρ*_*f*_ (*r, D*) scaled by *f* (*r, D*) is given by:

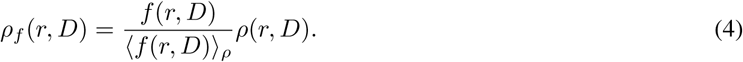

The function *f* (*r, D*) can be appropriately chosen so that the filtered PDF *ρ*_*f*_ (*r, D*) is able to provide specific information about the underlying joint distribution *ρ*(*r, D*). For example, if *f* (*r, D*) is a constant function over a fixed range of diffusivity, *D*_*a*_*< D < D*_*b*_, and zero elsewhere, i.e. an indicator function, then *ρ*_*f*_ (*r, D*) is equal to the conditional probability distribution *ρ*(*r, D | D*_*a*_*< D < D*_*b*_), which reflects the relaxation rates associated to a fixed range of diffusivities.

Based on the definition in Eq. (4), the moments of *ρ*_*f*_ (*r, D*) can be computed by

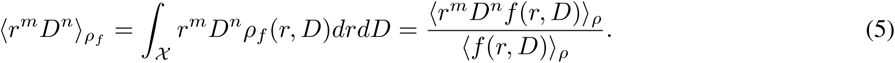

If *f* (*r, D*) is an analytic function of the form

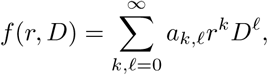

then

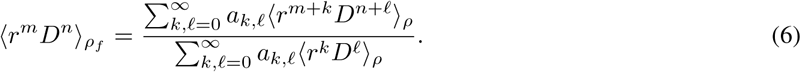

Since only the first three order moments of *ρ*(*r, D*) are estimated in Eq. (3), we restrict our analysis for *f*(*r, D*) using polynomials of order lower than three so that ⟨(*r*^*m*^*D*^*n*^*f* (*r, D*)⟩_*ρ*_ can be computed using the estimated parameters. Consequently, we consider two types of filters, o ne f or r escaling r elaxation, a nd o ne f or r escaling d iffusivity. For each type, we consider two filters t hat e mphasize s low a nd f ast r elaxation o r d iffusivity, r espectively, p roviding a total of four filters.

### Filters for different relaxation rates

We consider two filters that are able to emphasize dMRI signals from tissue components with slow and fast relaxation rates:

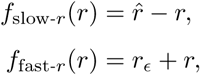

where *r*_*ϵ*_ is a positive constant to avoid singularity when computing ⟨*f*_fast-*r*_(*r, D*)⟩_*ρ*_. The two functions *f*_slow-*r*_ and *f*_fast-*r*_ impose relatively higher weight on signals from tissue components with slow and fast relaxation rates, respectively. If the underlying tissue components have equal relaxation rates, then the filtered joint PDFs 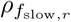 and 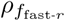 are equal to the original PDF *ρ*. On the other hand, if the underlying tissue components have heterogeneous relaxation rates, then the filtered P DFs are different from *ρ*(*r, D*), and these differences are reflected in their moments.

Using Eq. (6), we derive the following moments of the filtered j oint PDFs corresponding t o *f* _slow-*r*_ and *f*_fast-*r*_

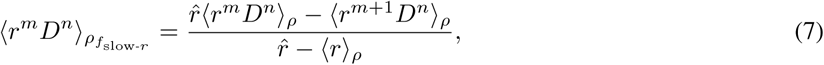

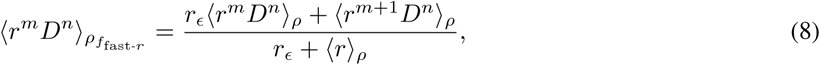

respectively. These moments can be applied to obtain diffusion signals contributed by tissue components with different relaxation rates, which will be explained in more detail later in this section.

### Filters for different diffusivities

Similarly, we define two types of multiplicative filters for characterizing hetero-geneous diffusion environments as

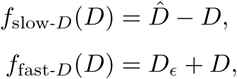

in order to emphasize tissue components with slow and fast diffusivities, respectively. The difference between the moments of the filtered PDFs, denoted by *ρ*_slow-*D*_ and *ρ*_fast-*D*_, reflect the underlying heterogeneity in diffusion coefficients. The moments of the filtered joint PDFs according to *f*_slow-*D*_ and *f*_fast-*D*_ are given by

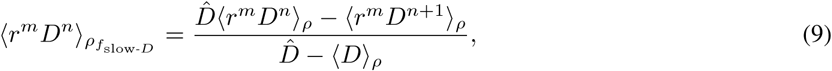

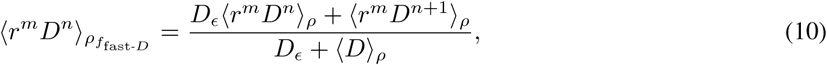

respectively. Next, we will use these moments to regress out the dependence of relaxation in diffusion signals.

### C. Marginal distribution of diffusivity and relaxation-regressed diffusion signal

The generative model in Eq. (1) implies that measured dMRI signals are weighted by the relaxation-related signal decay *e*^*−rt*^. Naturally, the estimated tissue microstructure and axonal orientations are also weighted by the echo time (TE). Thus the estimated tissue microstructure obtained using the standard dMRI may be biased by TE. This is especially true if the relaxation rates of crossing fiber bundles are different. The joint probability density function of relaxation and diffusion provides intrinsic information about the underlying diffusion process independent of TE. In particular, we define

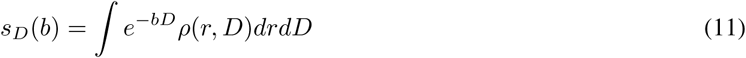

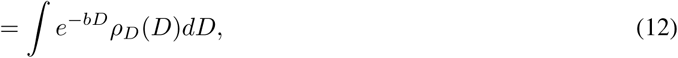

as the relaxation-regressed normalized diffusion signal where *ρ*_*D*_(*D*) = ∫ *ρ*(*r, D*)*dr* is the marginal PDF of diffusivity. The marginal PDF *ρ*_*D*_(*D*) contains information about the underlying diffusivity independent of TE which is not provided by the standard dMRI technique.

Based on the estimated moments, *s*_*D*_(*b*) can be approximated by the following cumulant expansion

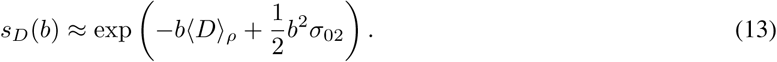

The kurtosis of *s*_*D*_(*b*) is given by

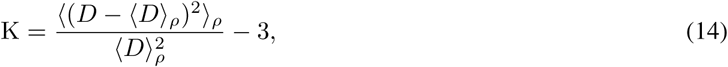

which reflects the non-Gaussianity of the probability distribution of water displacement independent of the T_2_-relaxation.

### D. Filtered relaxation-regressed diffusion signal

Similar to Eq. (11), we introduce the following family of relaxation-regressed dMRI signal based on the filtered joint PDFs:

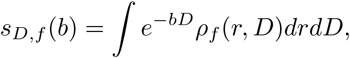

where *ρ*_*f*_ (*r, D*) denotes the filtered PDF using one of the multiplicative filters: *f*_slow-*r*_, *f*_fast-*r*_, *f*_slow-*D*_ and *f*_fast-*D*_. In particular, the filtered signals 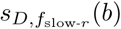 and 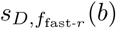 are scaled by the relaxation rate *r* instead of the TE-dependent weight function *e*^*−rt*^. On the other hand, 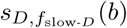 and 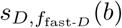 zoom into signals from components with slow and fast diffusivities, respectively.

Similar to (13), *s*_*D,f*_ (*b*) can be approximated by the cumulant expansion

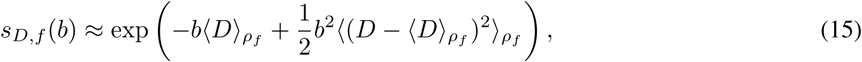

where the moments of the filtered PDFs can be obtained using Eqs. (6) to (10). The corresponding filtered kurtosis measures is equal to

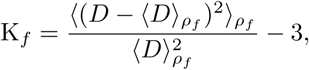

which provides more specific information on the probability distribution of displacements of water molecules in different tissue environments. As an example, the signal corresponding to the fast-r weighted filter is equal to

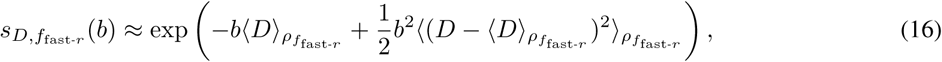

which can be computed using

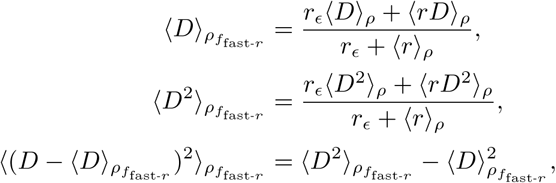

based on Eq. (8). The signal corresponding to other filters can be computed in a similar manner.

## III. METHODS

### A. Data and preprocessing

Diffusion MRI data from a healthy volunteer was acquired on a 3T Siemens Prisma scanner. The TE values were: 71, 101, 131, 161 and 191 ms. For each TE, dMRI images were acquired along 30 gradient directions at b = 700, 1400, 2100, 2800, 3500 s*/*mm^2^ together with 6 volumes at b=0. An additional pair of b=0 images with AP, PA phase encoding directions were also acquired along with a MPRAGE T1-weighted image. The voxel size of dMRI was 2.5 *×* 2.5 *×* 2.5 mm^3^. The size of data matrix is 96 *×* 96 *×* 54. The TR was fixed at 5900ms. Simultaneous multi-slice (SMS) technique with an acceleration factor of 2 was used along with iPAT=2. The total scan time was about 80 minutes.

The b0 volumes were corrected for EPI distortion correction by applying FSL TOPUP on reversed phase encoding pairs [21]. The rest of data was corrected for eddy current distortion, subject motion and EPI distortion with FSL TOPUP/eddy [22]. The MPRAGE T1-weighted images were used to create a FreeSurfer label map, which was subsequently rigidly registered to the distortion-corrected baseline volume using the ANTS toolbox [23].

### B. Analysis

We used Eq. (3) to estimate the moments in each voxel by solving a constrained least-squares problem using the Matlab function *lsqlin*. The constraints ensure that ⟨*r*⟩_*ρ*_, ⟨*D*(*u*))⟩_*ρ*_, *σ*_20_, *σ*_02_ are positive and ⟨*D*(*u*))⟩_*ρ*_*≤* 3 *µ*m^2^*/*ms. Based on the estimated moments, we used Eq. (6) to compute the moments of the filtered joint density of relaxation and diffusion where we fixed the following user-defined values for 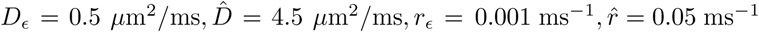. These values were chosen so that the filtered function *f* (*r, D*) is well-defined within the biophysical range of expected values.

Based on the estimated joint moments of relaxation and diffusion, we first compared the differences in fiber orientations between the relaxation-regressed signal and the measured data (TE-dependent). In particular, we applied the spherical-ridgelet-based method in [24] to estimate the fiber orientation distribution function (ODF) in a crossing-fiber region, as shown in Fig. 1. All ODFs were estimated using the diffusion signal at *b* = 1400 s*/*mm^2^ with the same regularization parameter *γ* = 0.005 that regularizes the sparseness of the representation coefficient in the basis of spherical ridgelet functions.

**Fig. 1:**
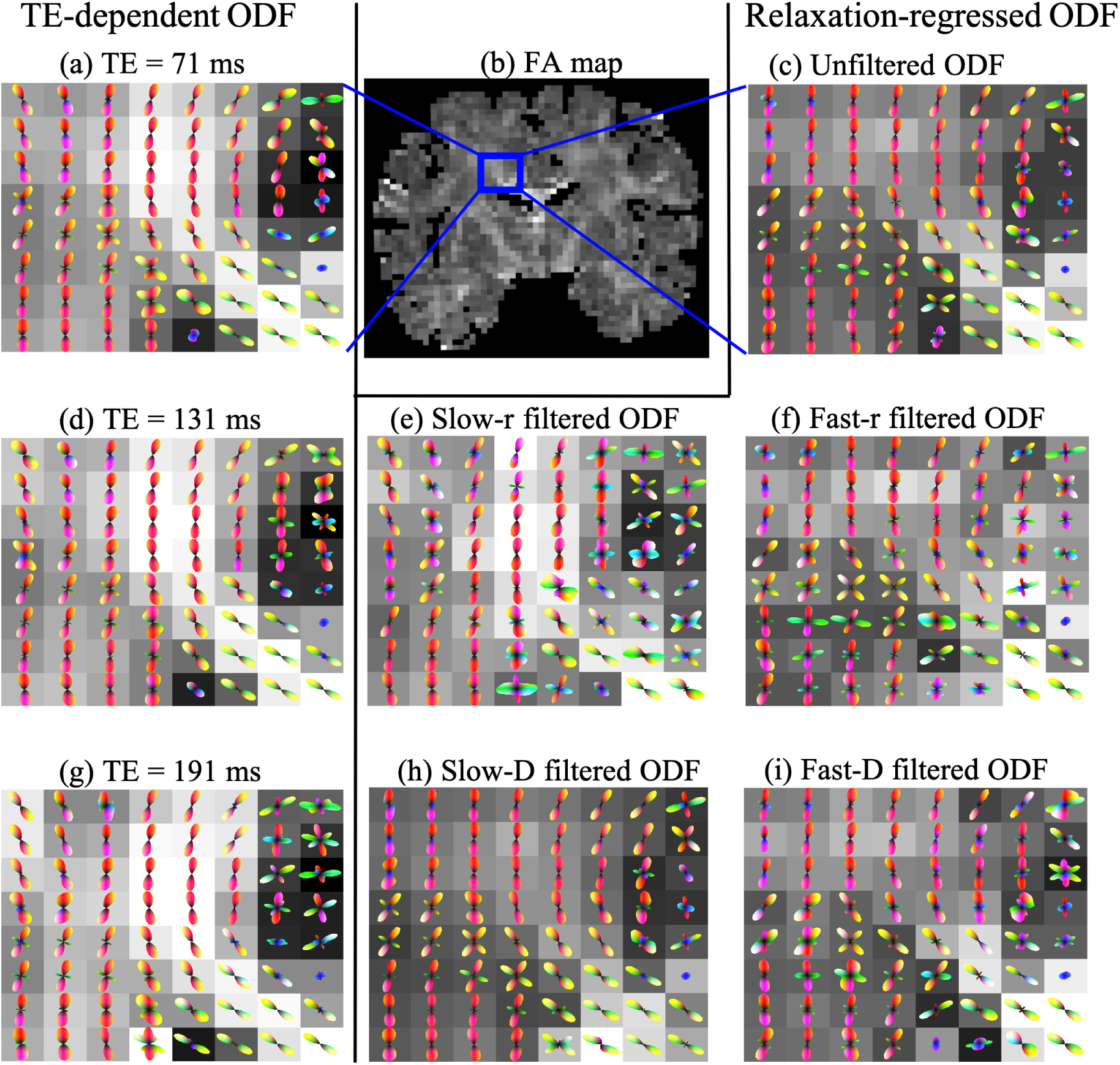
Comparisons between TE-dependent and relaxation-regressed orientation distribution functions (ODFs). Figs. (a), (d), (g) illustrate the ODFs based on the measured diffusion signals at TE= 71, 131, 191 ms within the brain region indicated by the box in (b). Figures (c), (e), (f), (h) and (i) illustrate the ODFs using the relaxation-regressed diffusion signals computed using Eq. (11) or Eq. (15) and the ODFs obtained using different filters.

We also computed several quantitative measures using the estimated joint moments. In particular, we computed the mean kurtosis (MK) [20]:

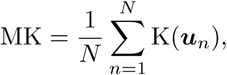

where K(***u***_*n*_) denotes the estimated kurtosis along gradient direction ***u***_*n*_ using Eq. (14), and *N* = 30 is the total number of gradient directions. Similarly, we also computed the average diffusivity:

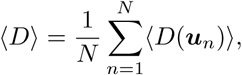

the mean diffusion-relaxation correlation coefficient:

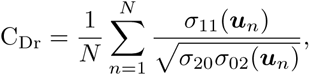

and the normalized variance:

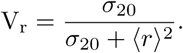

We note that the diffusivity computed above is not equal to the mean diffusivity from the diffusion tensor imaging model [25], though they are expected to have similar values.

From the joint moments computed using Eq. (5), we also computed MK, ⟨*D*⟩, C_Dr_ and V_r_ measures with respect to the filtered PDFs 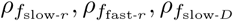 and 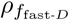. We further used the estimated moments in Eqs. (13) and (15) to compute the relaxation-regressed diffusion signal at *b* = 1400 ms*/µ*m^2^ along the same set of 30 gradient directions. Finally fractional anisotropy (FA) of the relaxation-regressed signal using the standard tensor model was also computed.

## IV. RESULTS

### I. A. Comparison of fiber orientations

An important result of this study shows that the crossing fiber bundles may have different (*T*_2_) relaxation rates. The left column of Fig. 1 shows the ODF in the centrum-semiovale and corpus callosum region indicated by the box in Fig. (5b) using the measured diffusion signal at TE = 71, 131, 191 ms, respectively. The left column of Fig. 2 shows the ODFs in four voxels with crossing fibers indicated by the box in Fig. (6b). It can be observed that the ODFs change with TE since fiber-bundles with faster relaxation rates contribute less to diffusion signals at long TE. As shown in Figs. (2a), (2d) and (2g), the volume fractions of different fiber bundles changes with TE. We believe this is the first time, this particular aspect of changes in volume fractions with changes in echo time has been probed and demonstrated.

**Fig. 2:**
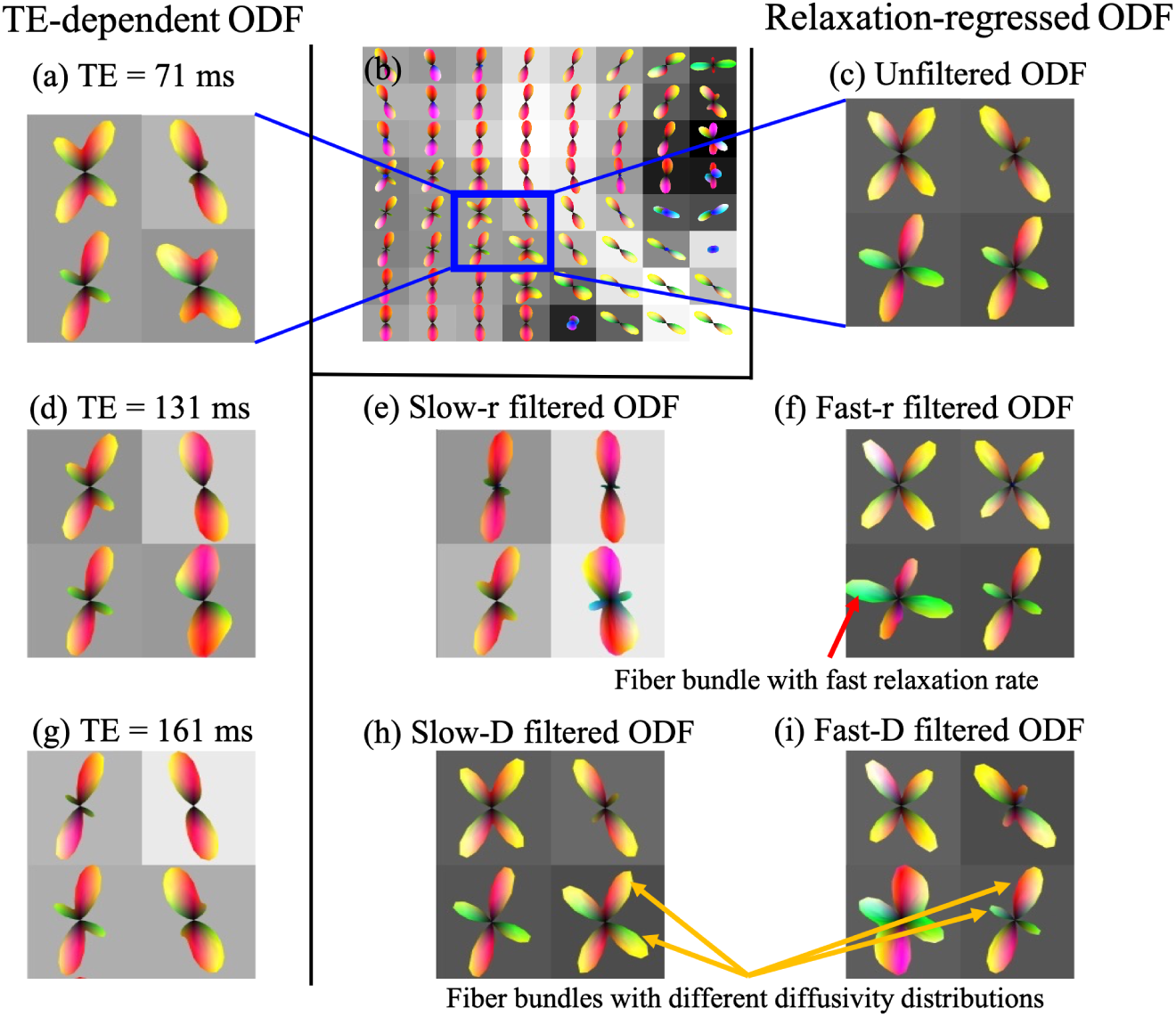
Comparison between TE-dependent and relaxation-regressed orientation distribution functions (ODFs) in four voxels with crossing fibers in the region indicated by the box in (b), which is the same as Fig. 1(a).

The right-hand side of Figs. 1 and 2 show the relaxation-regressed ODF based on the estimated joint PDFs. In particular, Figs. (1c) and (2c) show the ODF based on the marginal probability density *ρ*_*D*_(*D*) so that the diffusion signals are not scaled by T_2_ decays. As shown in the zoomed figures in Fig. (2c), the relative fraction of the crossing fibers and their orientations are all different from the TE-dependent ODF, which is potentially due to the effect of different transverse relaxation rates for different fiber bundles. Figs. (1e) and (2e) show the ODFs corresponding to slow-r weighted probability density function *ρ*_*D*,slow-*r*_(*D*) and Figs. (1f) and (2f) illustrate fast-r weighted ODFs. In these subplots (Figs. (1e) and (2e)), one can clearly observe only the the fiber ODFs corresponding to the corticospinal tract (Fig. (1e)), but not those originating from the corpus callosum. However, a contrasting scenario is seen for the fast-r weighted ODFs (Fig. (1f) and (2f)), where ODFs from the corpus callosum seem to dominate in the crossing fiber regions. This significant difference between the ODFs is because the corpus callosum fiber bundle in the crossing fiber region potentially has faster transverse relaxation rate compared to the corticospinal tract.

Figs. (1h), (1i) and Figs. (2h), (2i) illustrate slow-diffusion and fast-diffusion weighted ODFs, respectively. We note that the slow-diffusion weighted ODFs are very similar to the results in Figs. (1c) and (2c), implying that the underlying diffusion signals mostly come from tissue components with slow diffusion, e.g. the intra-axonal space. Compared to Figs. (2h), the ODFs in Fig. (2i) have blurred peaks and different volume fractions between the crossing fibers. These differences in Figs. (2h) and (2i) could potentially be due to the zooming effect into the fast diffusion component from the extra axonal space.

### B. Comparisons across brain regions

To appreciate the relation between the relaxation rate and diffusivity in various brain regions, in Figure 3, we show the contour plots of the relaxation rate and diffusivity in cortical-gray-matter, white-matter and sub-cortical-gray regions. The three rows illustrate the scatter plots corresponding to the standard (without filtering or weighting), diffusivity-weighted and relaxation weighted joint PDFs, respectively. The two-dimensional plots show the contour lines of the joint density between the relaxation and diffusivity values from all voxels within the ROIs shown in Fig. (3a). The one-dimensional plots on the top and right of the contour plots illustrate the marginal densities of relaxation and diffusion. Using the standard way to estimate the average diffusivity and relaxation in these regions (Figs. (3c) and (3d)) indicates the existence of a single type of homogeneous region in the white and sub-cortical gray matter region (the thalamus). For the cortical gray matter region in Fig. (3b), however, we observe a small multimodal peak for the relaxation component but with similar type of diffusion coefficient (unimodal distribution) throughout the region.

**Fig. 3:**
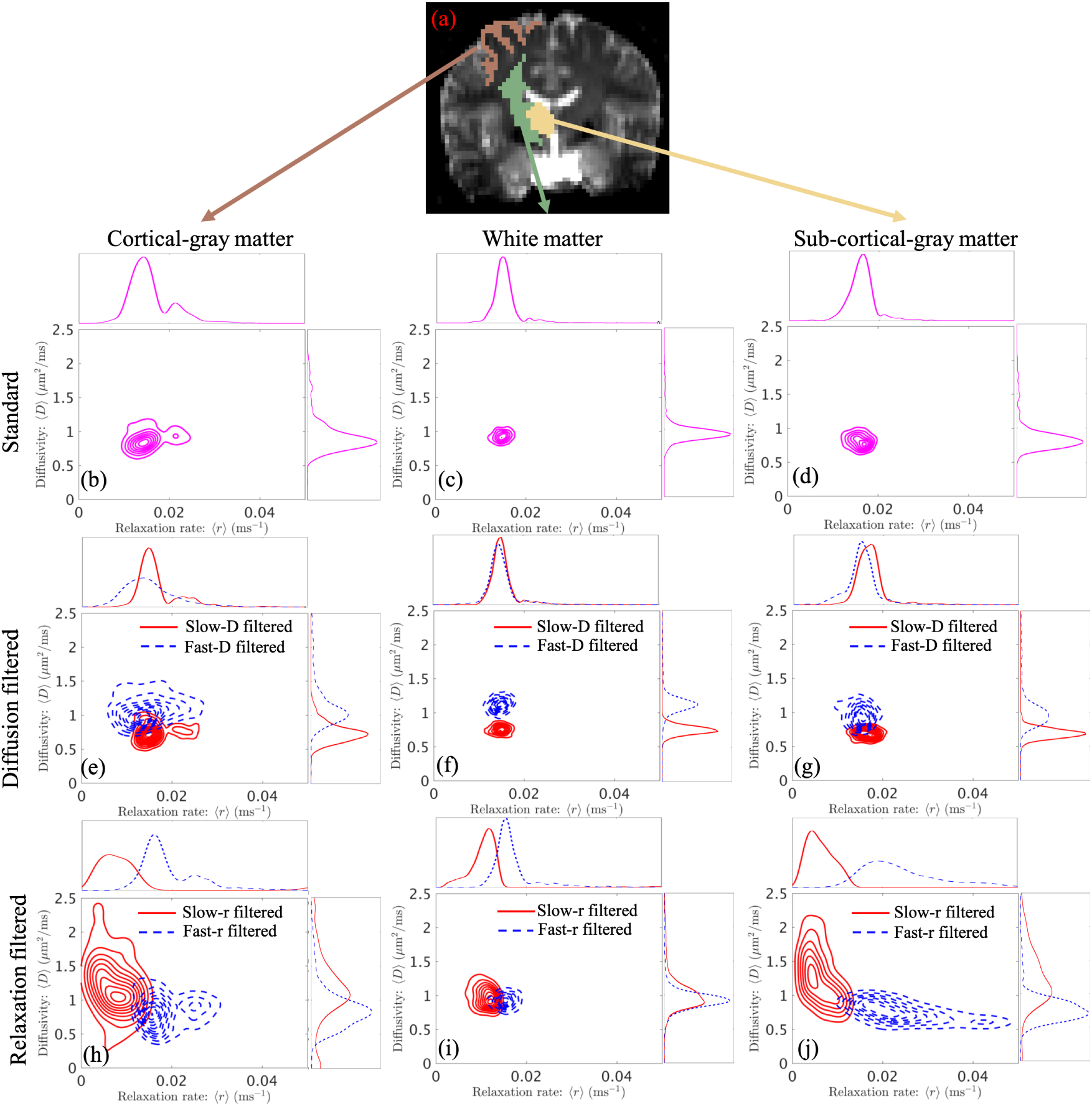
Contour plots of the joint distribution of relaxation rate and diffusivity in cortical-gray-matter (left), white-matter (middle) and subcortical-gray-matter (right) regions. The top row illustrates the scatter density of the standard joint results without any weighting functions. The middle row shows the scatter plots obtained using the proposed method corresponding to slow-diffusion and fast-diffusion weighted joint probability densities of relaxation and diffusion. The bottom row shows results for slow-and fast-relaxation filtered joint PDFs.

The results for filtered PDFs allow us to clearly visualize the heterogeneity of the tissue microstructure in all of the regions. For example, the average diffusivity has a multimodal distribution for the fast-D and slow-D weighted PDFs in all three regions (Figs. (3e), (3f) and (3g)). However, the relaxation rates for the fast-D and slow-D weighted PDFs in the white and sub-cortical gray mater regions are quite similar, with slight differences observed in the cortical regions.

As seen from Figs. (3h), (3i) and (3j), the average diffusivity in all three regions varies significantly with relaxation weighting, indicating high heterogeneity in the tissue composition. For example, for the cortical gray matter region (Fig. (3h)), we observe different relaxation and diffusivity for slow-r and fast-r filters. Similarly, we observe different values for the average relaxation and diffusivities for the filtered PDFs for both white and sub-cortical gray matter regions, which are not observed using the standard technique of estimating the moments (see Figs. (3c) and (3d)). The very similar results in slow-r and fast-r weighted densities in the white-matter region indicates that the underlying tissue components may have similar transverse relaxation rates. It is also interesting to note that the fast-r weighted density in the subcortical-gray-matter in Fig. (3j) spreads over regions close to *r* = 0.04 ms^*−*1^, which is equivalent to a T_2_ relaxation time of 25 ms. In standard dMRI, the value of TE is relatively long such that the MR signals from tissue components with very fast T_2_ relaxation rates, such as myelin, are more or less absent. However, the proposed fast-r weighted filters provide an approach to zoom into tissue components with fast relaxation rates that cannot be probed using current dMRI scans from clinical scanners.

### C. Comparisons across imaging measures

The first row of Figure 4 shows the mean relaxation rates corresponding to the standard and four types of weighted probability density functions. We observe significant differences between the slow-relaxation (slow-r) and fast-relaxation (fast-r) weighted results indicating heterogeneous relaxation rates at a voxel level in both white and gray matter. Similarly, the relaxation rate seems to be quite different between slow-diffusivity (slow-D) and fast-diffusivity (fast-D) weighted results.

**Fig. 4:**
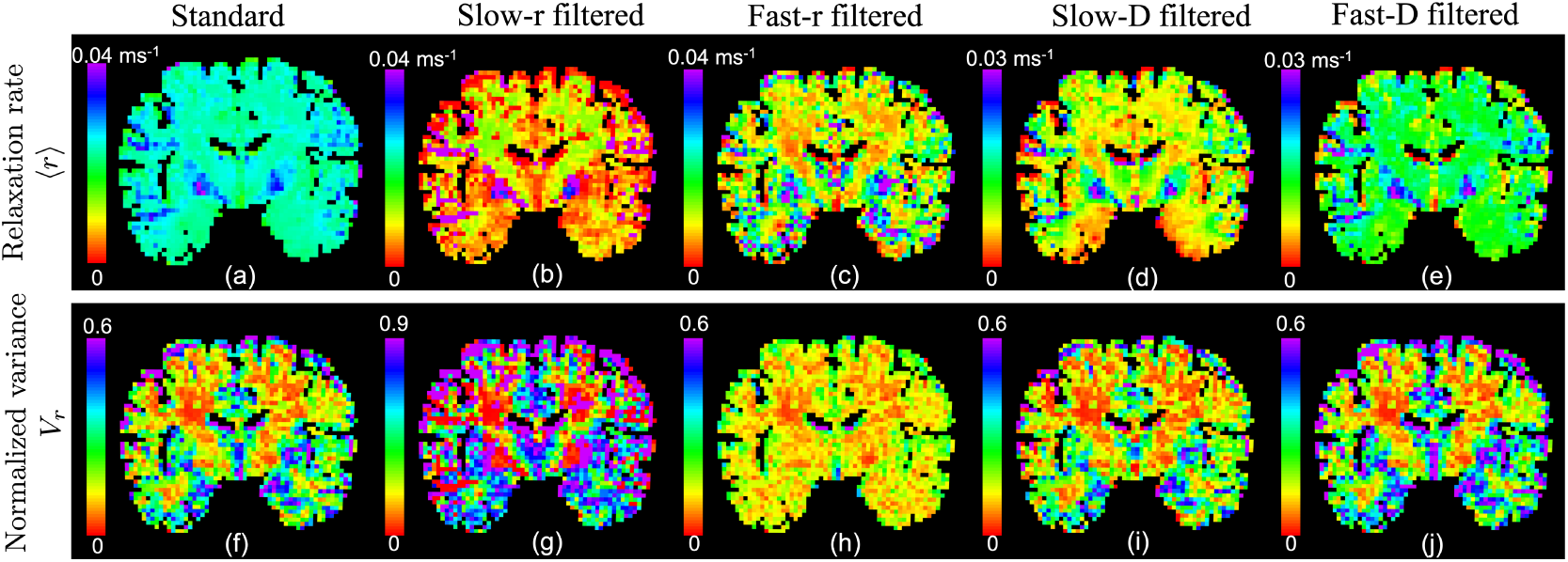
The first and second rows illustrate the mean and normalized variance of the relaxation rate corresponding to the standard and the four types of weighted joint probability density functions of relaxation and diffusion, respectively.

We note that in Fig. (4b) most cortical-gray matter regions have lower values than white-matter regions, whereas for the fast-r weighted result in Fig. (4c), there is an inverse contrast between the white and gray matter regions, implying that the underlying cortical-gray matter regions have much higher variance than the white matter regions. The slow-diffusivity weighted results show different contrast between cortical-gray matter, white matter and sub cortical gray matter regions. But the fast-D weighted results show similar values in most cortical-gray and white matter regions but much higher values in the subcortical gray matter regions.

The second row in Figure 4 shows the normalized variance of the relaxation rate. As expected, Fig. (2f) shows much higher variance in cortical-gray matter regions than in white matter regions. The higher variance in cortical gray matter regions is also consistently shown in Figs. (2g) to (2j). Moreover, Figs. (2i) and (2j) are very similar to each other whereas Figs. (2g) and (2f) are significantly different implying that the underlying tissue is potentially more heterogeneous in terms of the relaxation rates.

The three rows of Figure 5 illustrate the average diffusivity (⟨*D*⟩), the fractional anisotropy (FA) and the mean kurtosis (MK) as defined earlier in Section 3.2. We note that FA is computed based on dMRI signals at b = 1400 s*/*mm^2^. The first column shows the maps for these measures as estimated for the standard joint distribution *ρ*(*r, D*). For the slow-rsssssss weighted PDF 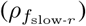, the diffusivity in cortical and subcortical gray-matter regions is much higher than in white-matter regions. But the fast-r weighted maps in the third column show slower diffusivity and higher kurtosis in gray-matter regions, implying that slow-r and fast-r weighted joint PDFs captures information from gray-matter tissue components with significantly different diffusion properties. The slow-r weighted PDF is potentially able to capture information from extra cellular spaces or large cells which act as free water components (and hence the slow relaxation rate), whereas fast-r weighted PDF contains information from intra-axonal or intra-cellular spaces.

**Fig. 5:**
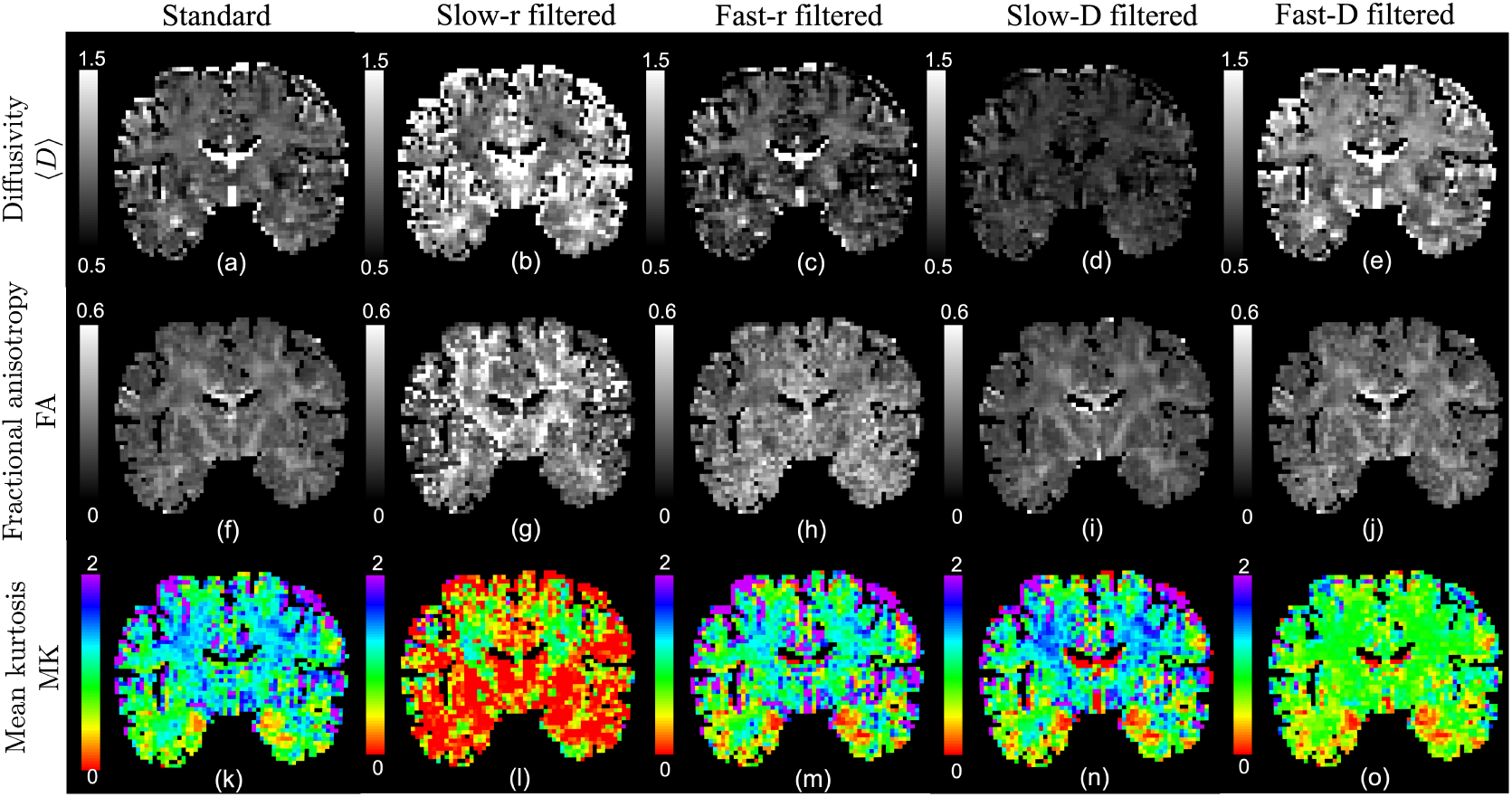
An illustration of the diffusivity ⟨*D*⟩, fractional anisotropy (FA) and mean kurtosis (MK) corresponding to the standard and the four types of weighted joint probability density functions of relaxation and diffusion.

The difference between slow-and fast-D weighted results show significant heterogeneity in the underlying diffusion. In particular, the slow-D weighted FA and MK maps are very similar to the standard results except for some voxels with potential partial-volume effects. This implies that the standard joint FA and MK measures are heavily weighted by the slow-D component. The similarity among fast-r and slow-D weighted results indicate that the slow-diffusion components potentially have fast relaxation rates, as expected.

Figure 6 shows the correlation coefficient *C*_*Dr*_ map between relaxation and diffusion. The results from the different joint PDFs have similar image contrasts in most regions except for some subcortical regions – see the circled regions. In these subcortical regions, the slow-r and fast-r weighted maps have negative and positive correlation, respectively, whereas in the slow-D, fast-D weighted as well as the standard results, the correlation is nearly zero. In all five maps, the cortical gray matter regions have significant negative correlation between diffusivity and relaxation rate.

**Fig. 6:**
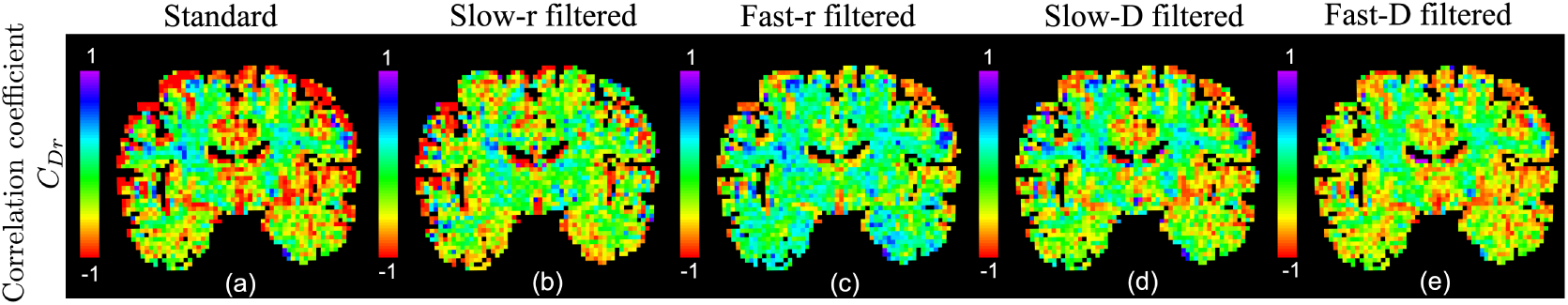
An illustration of the correlation coefficient between relaxation rate and diffusivity for different joint probability density functions.

## V. DISCUSSION AND CONCLUSIONS

In this work, we have introduced an approach to use the joint moments of relaxation and diffusion to derive multi-dimensional imaging measures and to compute relaxation-regressed diffusion signals for estimating fiber orientations. Our approach applied a multiplicative filter to the joint probability density function of relaxation and diffusion to zoom into the statistical properties of the different types of tissue components. We also introduced an expression for the joint moments of the filtered PDFs in terms of the original joint moments of relaxation and diffusion. Theoretically, any type of filter can be applied to select specific diffusion or relaxation bands in the underlying joint distribution. However, for more practical purposes of use in clinical settings and the lack of reliability of the estimated high-order moments from in-vivo clinical data, we introduced four types of linear filters to zoom into tissue components with slow relaxation, fast relaxation, slow diffusion and fast diffusion.

In a proof-of-concept experiment, we applied the proposed method to analyze a dataset acquired from a 3T Siemens Prisma scanner. Experimental results showed that the filtered joint PDFs of relaxation and diffusion had significant differences in their moments, implying heterogeneous components in the underlying tissue. In particular, cortical- and subcortical-gray-matter regions showed much higher variances in relaxation and more significant negative correlation coefficients between relaxation and diffusion than white matter region. The results also revealed different relaxation rates between axonal bundles in complex white matter regions. For the first time, using the proposed relaxation-regressed diffusion signals with different filters, we were able to separate two crossing fiber bundles with different transverse relaxation rates.

A limitation of the proposed method is still the requirement for relatively long scan time. Though the least-squares algorithm is expected to be more reliable than inverse-Laplace transform based methods using fewer number of samples, the in-vivo dataset used in the examples still required 90 minutes of scan time. Our future work will focus on optimizing data acquisition and estimation methods in order to reduce scan time and make this approach practical in clinical settings.

To summarize, the proposed method has several favorable features. First, the joint moments can be easily estimated compared to the ill-posed estimation of the full joint probability density function. The method is also more robust compared to multi-compartment models, which is plagued by non-unique solutions. Second, the standard non-parametric approach for estimating fiber orientations can be applied to relaxation-regressed diffusion signals to compute the orientation of different fiber components. Third, the imaging measures and fiber orientations from different type of filters could potentially be more sensitive to brain abnormalities than the standard diffusion data acquired at a fixed TE. Therefore, the proposed method provides a new direction to investigate brain structural connections and estimation of their microstructure.

